# Cleavage of MALAT1 RNA by 14-nt sgRNA-guided tRNase Z^L^

**DOI:** 10.1101/2025.01.28.635233

**Authors:** Masayuki Takahashi, Masayuki Nashimoto

## Abstract

We have been developing a gene suppression technology, tRNase Z^L^-utilizing efficacious (TRUE) gene silencing, in which artificially designed small guide RNA (sgRNA) guides tRNase Z^L^ to cleave cellular target RNA. In this study, we examined 14-nt linear-type sgRNAs, which are fully 2′-*O*-methylated and have full phosphorothioate linkages, for their ability to suppress a level of a nuclear-localized long non-coding RNA, Metastasis Associated Lung Adenocarcinoma Transcript 1 (MALAT1). The MALAT1 RNA is implied to be involved in stress responses and diseases including cancers. Specifically, we designed six 14-nt linear-type sgRNAs, sgRM1−sgRM6 that target the human MALAT1 RNA. sgRM1, sgRM2 and sgRM6 suppressed the MALAT1 RNA level, while the other sgRNAs showed little effect. In order to demonstrate that the suppression effect of sgRM1, sgRM2 and sgRM6 on the MALAT1 RNA level is caused by TRUE gene silencing, we performed *in vitro* tRNase Z^L^ cleavage assay, microscopic analysis for nuclear existence of sgRNA, and tRNase Z^L^ knockdown experiment. For the *in vitro* tRNase Z^L^ cleavage assay, three 30-nt MALAT1 RNA fragments, TM1, TM2 and TM6 were prepared, which were RNA targets for sgRM1, sgRM2 and sgRM6, respectively. All of the sgRNAs guided recombinant tRNase Z^L^ *in vitro* to cleave their own targets, although the cleavage efficiency changed depending on target/sgRNA pairs. By fluorescence microscopy, a 14-nt 5′-Alexa568-labeled sgRNA released from liposome was observed to be distributed ubiquitously in A549 cells with higher density in the nucleus, where both the target MALAT1 RNA and tRNase Z^L^ exist. Knockdown of tRNase Z^L^ by siRNA attenuated the suppression effect of sgRM1, sgRM2 and sgRM6 on the MALAT1 RNA level. We also demonstrated that the effective sgRNAs sgRM1, sgRM2 and sgRM6 reduce A549 cell viability.

## Introduction

The discovery in 1991 of a sequence-specific endoribonuclease activity that needs a cellular tRNA fragment led us to attempt to develop a gene suppression technology, tRNase Z^L^-utilizing efficacious (TRUE) gene silencing [1-3]. Although appearing to have a clinical potential, this technology has been progressing slowly and is still in its infancy [4-6]. TRUE gene silencing is based on the property of tRNase Z^L^ that it can recognize and cleave a pre-tRNA-like or micro-pre-tRNA-like complex [3]. Such a complex can be formed between a target RNA and an artificially designed small guide RNA (sgRNA), which is categorized into four groups, 5′-half-tRNA-type, hook-type, ∼14-nt linear-type, and heptamer-type.

We have been attempting to apply TRUE gene silencing to multiple myeloma or leukemia to find a cure by searching for effective sgRNAs [7-12]. In this attempt, among the four types, heptamer-type sgRNAs were chosen, which were 5′-/3′-phosphorylated and fully 2′-*O*-methylated, considering cost performance, nuclease resistance, and the efficiency in guiding tRNase Z^L^ [13]. We have recently, however, shown that longer oligonucleotides are intracellularly taken up more efficiently and that the oligonucleotides with phosphorothioate linkages are taken up by cells more efficiently than those without the linkages [14]. Taking account of these observations, here, we examined 14-nt linear-type sgRNAs, which are fully 2′-*O*-methylated and have full phosphorothioate linkages, for their ability to suppress a level of a nuclear-localized long non-coding RNA (lncRNA), Metastasis Associated Lung Adenocarcinoma Transcript 1 (MALAT1). The MALAT1 RNA is implied to be involved in stress responses and diseases including cancers [15]. And downregulation of the MALAT1 RNA level by antisense oligonucleotides (ASOs) that can guide RNase H1 has been shown to suppress metastasis of human lung cancer cells or growth of human myeloma cells in a mouse xenograft model [16,17].

## Materials and Methods

### Oligonucleotide preparation

The following oligoribonucleotides were chemically synthesized and subsequently purified by high-performance liquid chromatography by Nippon Bioservice (Saitama, Japan): six 5′-/3′-phosphorylated, fully 2′-*O*-methylated sgRNAs with full phosphorothioate modifications, sgRM1 (5′-UAGGAUUCUAGACA-3′), sgRM2 (5′-UGGUUAUGACUCAG-3′), sgRM3 (5′-AUUGCCUCUUCAUU-3′), sgRM4 (5′-CCUUCUGCCUUAGU-3′), sgRM5 (5′-CUUUUGCAUUUCCC-3′), and sgRM6 (5′-AAUCCCCUAGGGAA-3′), and a 5′-Alexa568-labeled, 3′-phosphorylated, fully 2′-*O*-methylated sgRNA with full phosphorothioate linkages (5′-UUGGUACCUUCUCA-3′).

The following primers used for quantitative reverse-transcription polymerase chain reaction (qRT-PCR) were obtained from Fasmac (Kanagawa, Japan): MALAT1 forward primer, 5′-GGATCCTAGACCAGCATGCC-3′; MALAT1 reverse primer, 5′-AAAGGTTACCATAAGTAAGTTCCAGAAAA-3′; β-actin forward primer, 5′- ACAATGTGGCCGAGGACTTT-3′; β-actin reverse primer, 5′- TGTGTGGACTTGGGAGAGGA-3′.

Three 5′-FAM labeled MALAT1 RNA fragments, TM1 (5′- GGUCUGUCUAGAAUCCUAAAGGCAAAUGAC-3′), TM2 (5′- UCUUCUGAGUCAUAACCAGCCUGGCAGUAU-3′), and TM6 (5′- CCCCUUCCCUAGGGGAUUUCAGGAUUGAGA-3′) were chemically synthesized and subsequently purified by high-performance liquid chromatography by Ajinomoto Bio-Pharma (Tokyo, Japan).

siRNAs, sitRNase Z^L^ (sense strand sequence: 5′- CGCUGUUGCGAACAUGUGAUU-3′) and siControl (sense strand sequence: 5′- CAGCACGACUUCUUCAAGUCC-3′) were obtained from Nippon Bioservice (Saitama, Japan).

### Cell culture

Human A549 cells [18] were cultured in DMEM (Wako, Osaka, Japan) supplemented with 10% fetal bovine serum (COSMO BIO, Tokyo, Japan) and at 37°C in a humidified incubator with 5% CO2.

### Transfection

Transfection was performed using Lipofectamine 3000 (Thermo Fisher Scientific, Tokyo, Japan) following the manufacturer’s protocol for reverse transfection. In tRNase Z^L^ knockdown experiments, A549 cells were transfected with 20 nM siRNA, and after 24-hr culture, they were transfected with 200 nM sgRNA and further cultured for 96 hr.

### qRT-PCR

Total RNA was extracted from A549 cells using TRI Reagent (Molecular Research Center, Cincinnati, USA), and cDNA was synthesized using PrimeScript RT Reagent Kit (Perfect Real Time) (Takara, Shiga, Japan) according to the manufacturer’s protocol. MALAT1 RNA, tRNase Z^L^ mRNA and β-actin mRNA were quantified by PCR with THUNDERBIRD SYBR qPCR Mix (TOYOBO, Osaka, Japan) using a Thermal Cycler Dice Real Time System (Takara).

### Preparation of recombinant human tRNase Z^L^

The histidine-tagged human tRNase Z^L^ that lacks N-terminal 30 amino acids was overexpressed using the expression plasmid pQE-80L in *E. coli* strain Rosetta(DE3)pLysS and purified with nickel-agarose beads as reported previously [19].

### *In vitro* tRNase Z^L^ cleavage assay

*In vitro* tRNase Z^L^ cleavage assay was performed at 37°C with 1 pmol of 5′-FAM- labeled target RNA, 1 pmol of sgRNA and 8 pmol of recombinant human tRNase Z^L^ in a 20-μl mixture containing 10 mM Tris–HCl (pH 7.1) and 10 mM MgCl_2_. Reaction products were resolved on a 12.5% polyacrylamide gel containing 8 M urea and analyzed using a Gel Doc EX imager (Bio-Rad, Tokyo, Japan).

### Fluorescence microscopy

A549 cells were prepared in a 35-mm glass-bottom dish (IWAKI, Tokyo, Japan). Forty-eight hr after transfection with the 14-nt Alexa568-labeled sgRNA, the cells were fixed with 4% paraformaldehyde in phosphate-buffered saline (PBS) for 15 min, permeabilized with 0.5% Triton X-100 in PBS for 15 min, and incubated with primary antibodies against a human tRNase Z^L^ peptide (amino acid 812–826) for 1 hour.

Subsequently, the cells were incubated with an Alexa488-conjugated secondary antibody (COSMO BIO) for 1 hr. Hoechst 33342 (Dojindo, Kumamoto, Japan) was used for nuclear staining. Fluorescence imaging was performed using an All-in-One Fluorescence Microscope BZ-X800 (KEYENCE, Osaka, Japan).

### Western blotting

A549 cells were lysed with a buffer containing 50 mM Tris-HCl (pH 7.5), 1% Triton- X100, 150 mM NaCl, and 5 mM EDTA, and the lysates were separated on an SDS/7.5% polyacrylamide gel, and transferred to a nitrocellulose membrane. tRNase Z^L^ and β-actin on the membrane were detected by the rabbit polyclonal antibodies raised against the human tRNase Z^L^ peptide (amino acid 812–826) and a mouse monoclonal antibody against human β-actin (Sigma-Aldrich, Tokyo, Japan), respectively, together with an anti-rabbit IgG HRP conjugated secondary antibody and an anti-mouse IgG HRP conjugated (R&D systems, Minneapolis, USA) using a LAS-3000 (FUJIFILM, Tokyo, Japan).

### Cell viability assay

A549 cells were plated at a density of 80,000 cells per well in 500 μL of medium on a 24-well dish, and transfected with 200 nM of sgRNA. After 96-hr culture, cell viable was measured using Cell Counting Kit-8 (Dojindo).

## Results

### Six 14-nt linear-type sgRNAs that target the human MALAT1 RNA

We would have a better chance to suppress the MALAT1 RNA level by TRUE gene silencing if we can select its target sites in less folded regions. However, since we do not know accurate tertiary structures of the MALAT1 RNA in human cells, we chose three sites that have been reported to be good target sites for ASOs. The 14-nt linear-type sgRNAs for the MALAT1 RNA sgRM1 and sgRM2 (Fig 1) were designed at the sites where two ASOs have been demonstrated to effectively suppress the MALAT1 RNA level in human EBC-1 cells [16]. And the 14-nt linear-type sgRNA sgRM3 (Fig 1) was designed at the site where an ASO has been shown to effectively reduce the MALAT1 level in human myeloma cells [17]. In addition, three 14-nt linear-type sgRNAs sgRM4−sgRM6 (Fig 1) were designed at arbitrarily selected sites. These sgRNAs were chemically synthesized as 5′-/3′-phosphorylated and fully 2′-*O*- methylated oligonucleotides with full phosphorothioate linkages.

**Fig 1.**
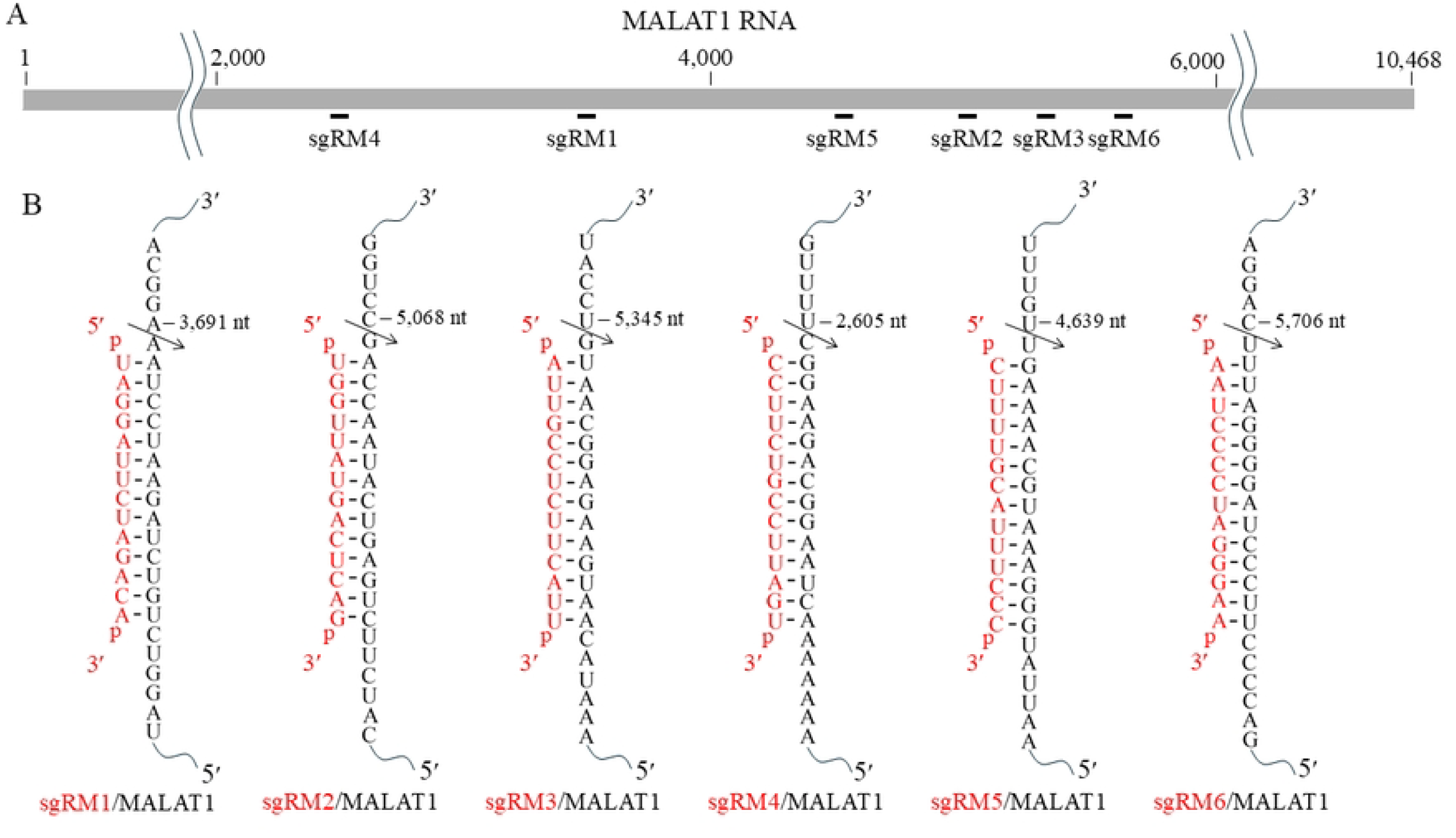
Six 14-nt sgRNAs and their target, human MALAT1 RNA. (A,B) Rough binding positions of the sgRNAs sgRM1−sgRM6 on the MALAT1 RNA (A) and sequences of the target/sgRNA complexes (B) are shown. An arrow represents an expected cleavage site.

### Suppression of the MALAT1 RNA level by 14-nt sgRNAs

We examined the six sgRNAs for their effect on the MALAT1 RNA level in human A549 cells. The cells were transfected with 200 nM of each sgRNA and cultured for 24 −96 hr, and the MALAT1 RNA in the cells was quantitated by qRT-PCR. While all the sgRNAs showed little effect on the MALAT1 RNA level in 24 and 48 hr, its amounts were reduced in 96 hr by sgRM1, sgRM2 and sgRM6 to 42, 68 and 68%, respectively (Fig 2A).

**Fig 2.**
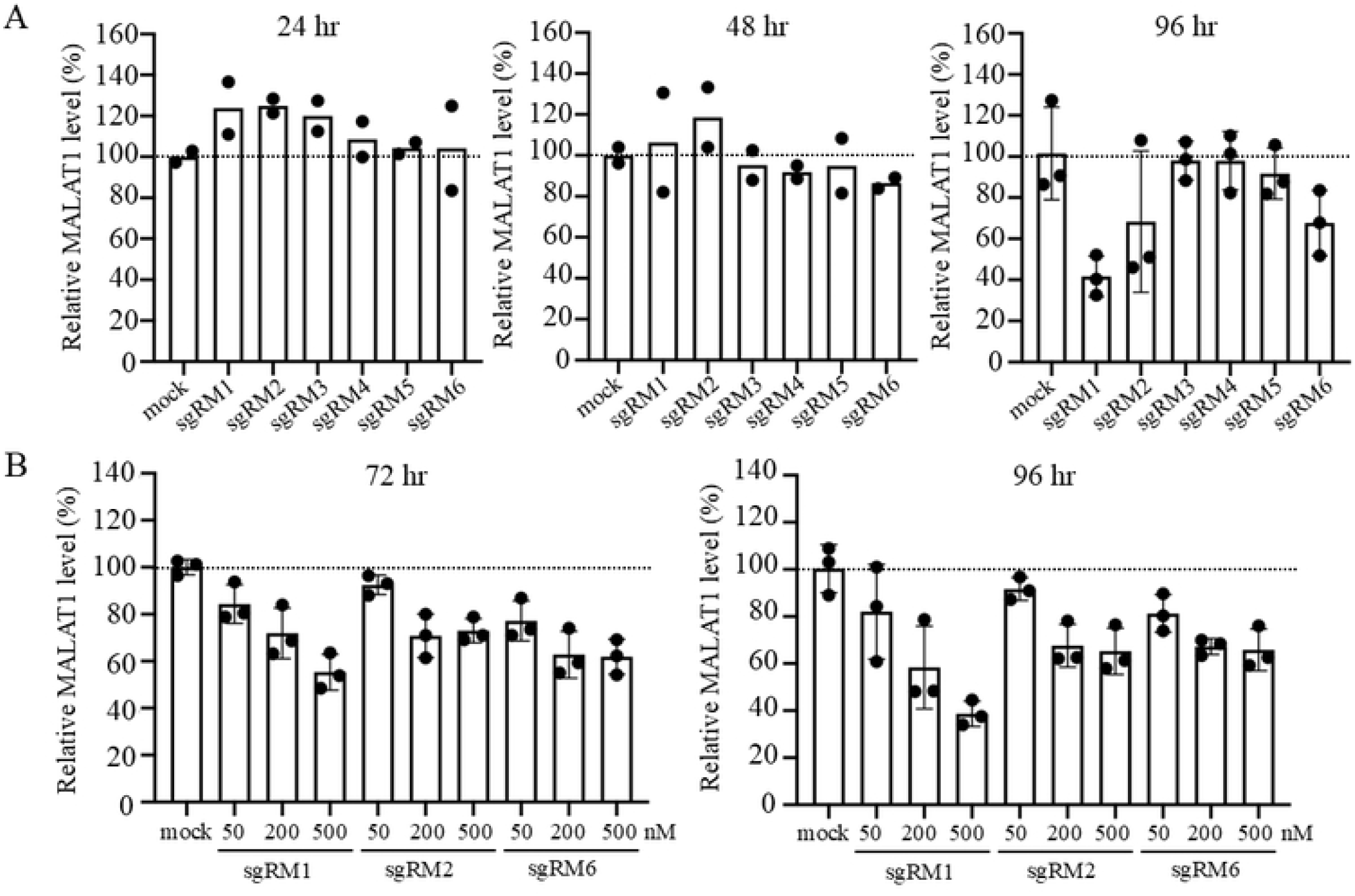
Suppression of a human MALAT1 RNA level by 14-nt linear-type sgRNAs. (A) Human A549 cells were transfected without (mock) and with each of the sgRNA sgRM1−sgRM6 (200 nM). After 24-, 48- and 96-hr culture, a MALAT1 RNA amount was measured by qRT-PCR, normalized against a β-actin mRNA amount, and expressed as a percentage relative to that of untreated cells. Values are mean for two biological replicates or mean ± SD for three biological replicates. (B) The A549 cells were transfected without (mock) and with 50, 200 and 500 nM of each sgRNA, sgRM1, sgRM2 or sgRM6. After 72- and 96-hr culture, a MALAT1 RNA level was similarly analyzed. Values are mean ± SD for three biological replicates.

With respect to sgRM1, sgRM2 and sgRM6, we analyzed dose dependency of their effect. The analysis for 50, 200 and 500 nM of sgRNA was carried out in 72 and 96 hr. Dose dependency of sgRM1’s effect was clearly shown, whereas the effect of sgRM2 and sgRM6 seemed to be saturated over 200 nM (Fig 2B).

### The 14-nt sgRNAs can guide tRNase Z^L^ to cleave the MALAT1 RNA

In order to demonstrate that the suppression effect of sgRM1, sgRM2 and sgRM6 on the MALAT1 RNA level is caused by TRUE gene silencing, we performed *in vitro* tRNase Z^L^ cleavage assay, microscopic analysis for nuclear existence of sgRNA, and tRNase Z^L^ knockdown experiment.

For the *in vitro* tRNase Z^L^ cleavage assay, we prepared three 30-nt MALAT1 RNA fragments, TM1, TM2 and TM6, which were 5′-FAM labeled RNA targets for sgRM1, sgRM2 and sgRM6, respectively (Fig 3). These 14-nt sgRNAs were tested for their ability to guide recombinant tRNase Z^L^ to cleave their MALAT1 targets. The cleavage reaction was carried out at 37°C for 15, 30 or 60 min, and the percent cleavage was graphed (Fig 3). All of the sgRNAs guided tRNase Z^L^ *in vitro* to cleave their own targets, although the cleavage efficiency changed depending on target/sgRNA pairs.

**Fig 3.**
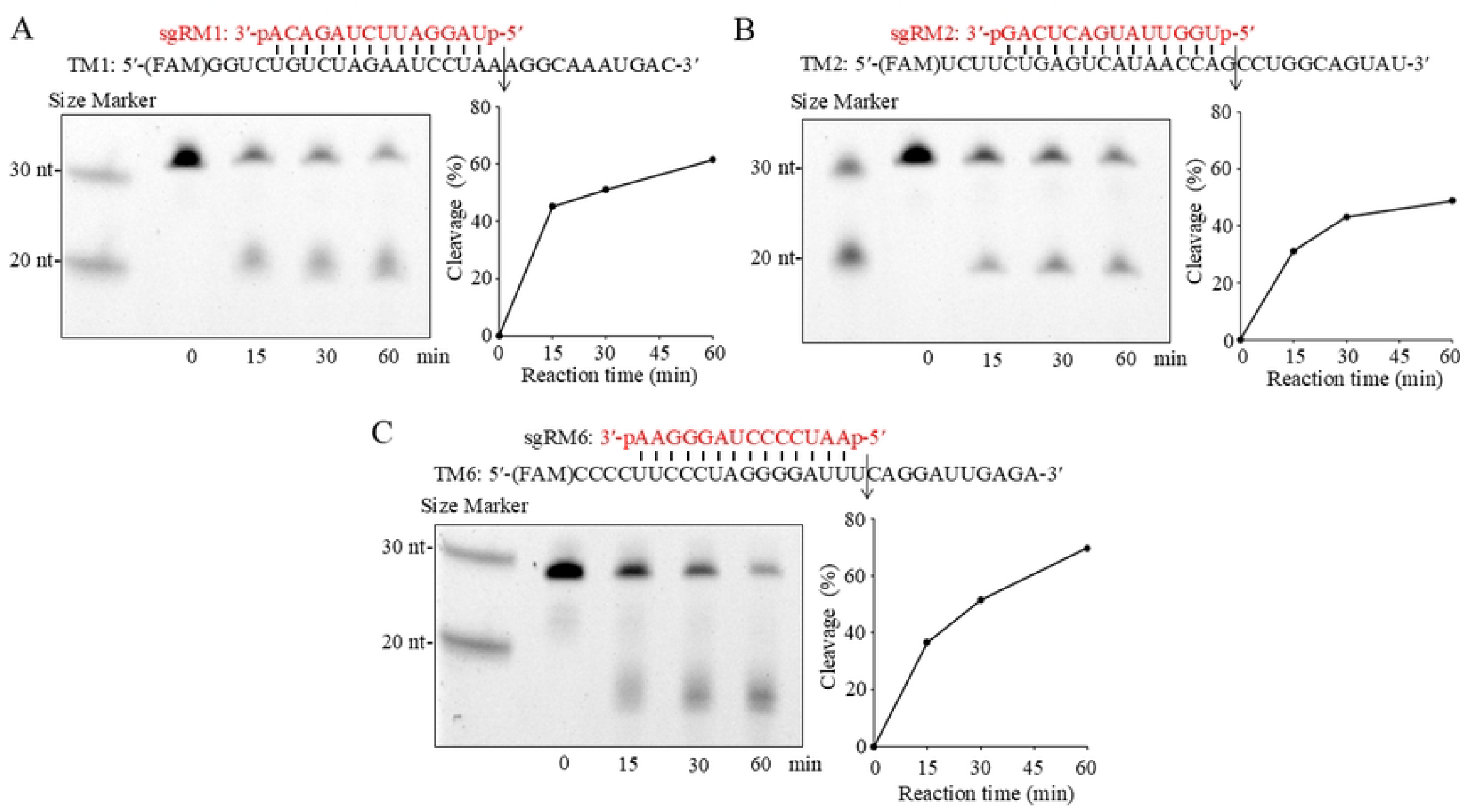
*In vitro* tRNase Z^L^ cleavage assay. The 30-nt 5′-FAM-labeled MALAT1 fragments TM1, TM2 and TM6 were incubated in the presence of sgRM1, sgRM2 and sgRM6, respectively, with recombinant human tRNase Z^L^ at 37°C for 0, 15, 30 and 60 min. A cleavage product was analyzed on a denaturing 12.5% polyacrylamide gel, and percent cleavage is graphed. An arrow denotes an expected cleavage site.

The targets TM1 and TM2 were cleaved by the corresponding sgRNA-guided tRNase Z^L^ at the expected sites, although the exact cleavage sites were not determined (Fig 3A,B). Likewise, the target TM6 was cleaved by sgRM6-guided tRNase Z^L^ (Fig 3C). The reason that the target TM6 and its 5′ cleavage product migrated on a polyacrylamide gel slightly faster than expected may be due to hairpin formation through base-pairing between a 4-nt cytosine stretch and a 4-nt guanine stretch in TM6.

### A 14-nt sgRNA encapsulated into liposome can be delivered to the nucleus

Intracellular distribution of sgRNA released from liposome was analyzed with a 14-nt test sgRNA, which was a 5′-Alexa568-labeled, 3′-phosphorylated and fully 2′-*O*-methylated oligonucleotide with full phosphorothioate linkages. As in Fig 4, the sgRNA was observed to be distributed ubiquitously in A549 cells 48 hr after transfection, with higher density in the nucleus.

**Fig 4.**
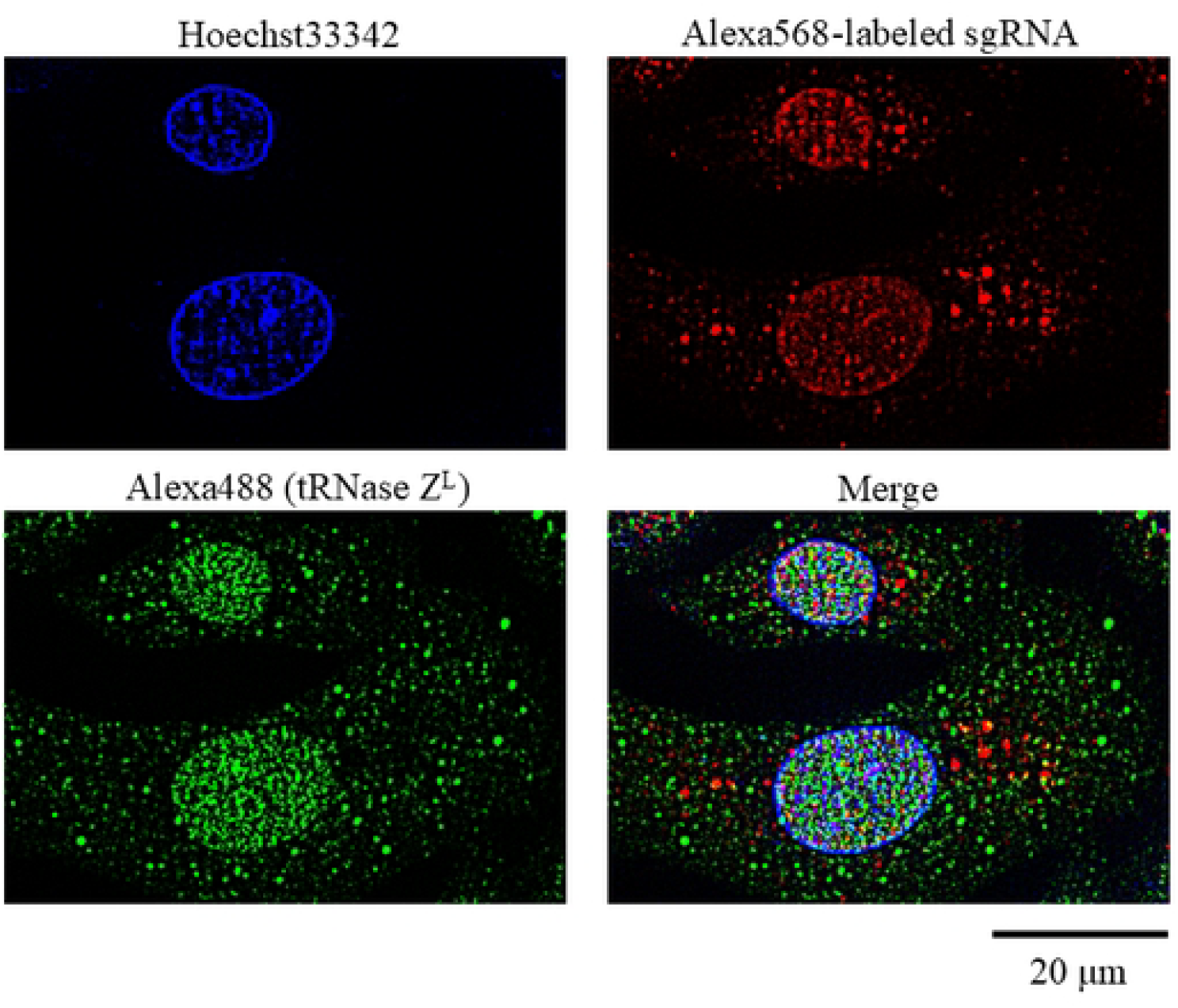
Subcellular distribution of a 14-nt linear-type sgRNA and tRNase Z^L^ in A549 cells. A549 cells were transfected with a 14-nt 5′-Alexa568-labeled linear-type sgRNA, and after 48-hr culture, the cells were analyzed with a fluorescence microscope. tRNase Z^L^ was visualized with an Alexa488-conjugated secondary antibody. Hoechst 33342 was used to stain the nucleus.

It has been reported that tRNase Z^L^ is distributed ubiquitously in various human cells, although the subcellular density distribution appears to differ depending on cellular conditions and/or analytical conditions [19−21]. In this study, tRNase Z^L^ in A549 cells was observed to be distributed ubiquitously with much higher density in the nucleus (Fig 4). Considering nuclear localization of the MALAT1 RNA, the observation that sgRNA encapsulated into liposome can be delivered easily to the nucleus would provide a rationale for suppression of the MALAT1 RNA level by TRUE gene silencing.

### tRNase Z^L^ is responsible for suppression of the MALAT1 RNA level

If tRNase Z^L^ is responsible for the suppression of the MALAT1 RNA level shown in Fig 2, downregulation of tRNase Z^L^ would attenuate the suppression effect. To examine this attenuation, we analyzed a MALAT1 RNA level in A549 cells, where a tRNase Z^L^ amount was reduced using siRNA (Fig 5A). The MALAT1 level in the presence of sgRM1 was increased by 56% (from 27 to 42%) in the tRNase Z^L^ knockdown cells compared with that in the normal cells (Fig 5B). This assay was repeated twice, and levels of the attenuation effect by tRNase Z^L^ knockdown were by 51 and 28% (Fig 5C,D). Similarly, the attenuation effect by tRNase Z^L^ knockdown was observed for the cells transfected with sgRM2 and sgRM6, in which levels of the attenuation effect were by 82−98% and by 25−38%, respectively (Fig 5B-D). Reduction of the tRNase Z^L^ level in the cells transfected with the ineffective sgRM5 (Fig 2A) showed only a trivial effect (Fig 5B-D).

**Fig 5.**
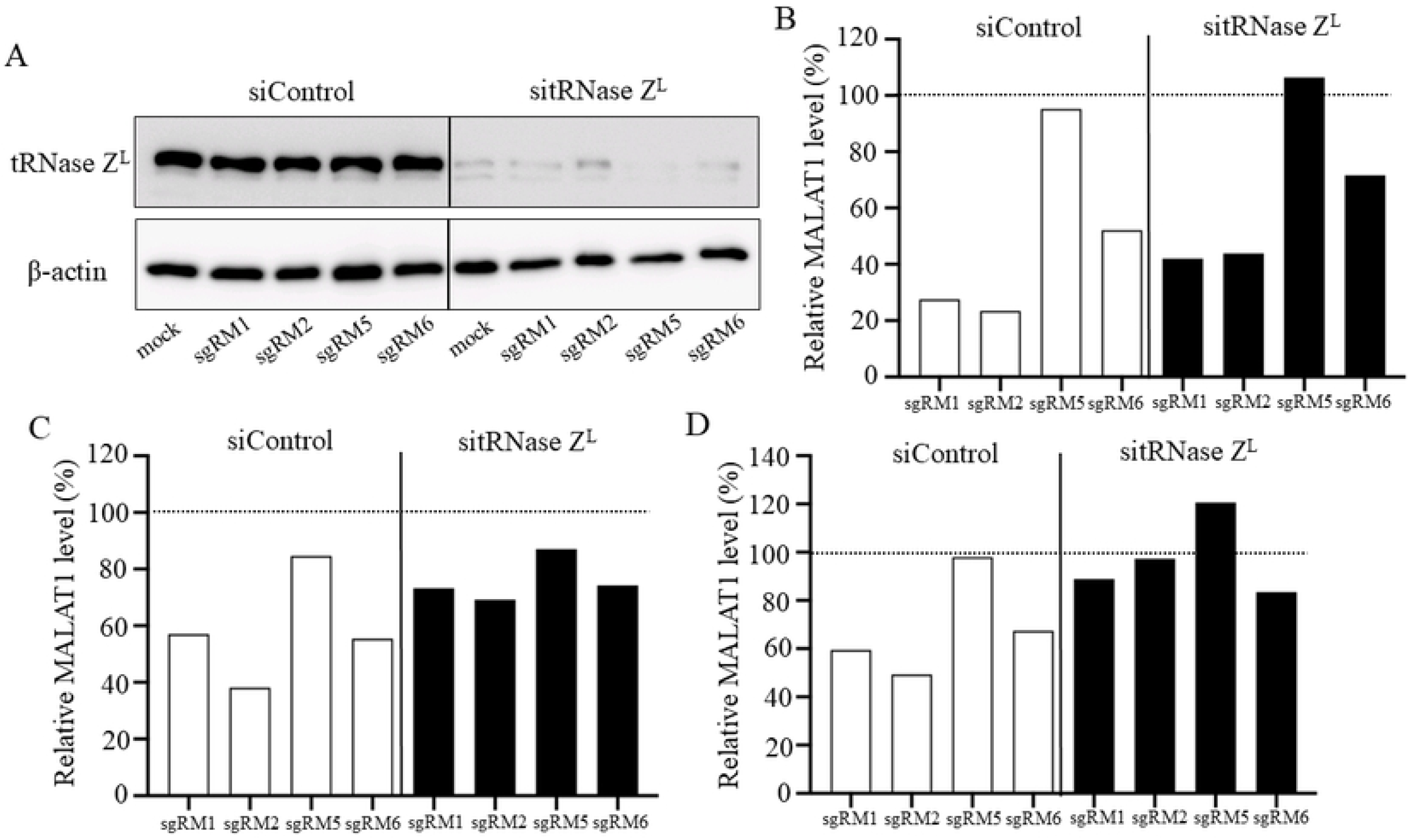
Knockdown of tRNase Z^L^ attenuates the suppression effect of sgRNA on the MALAT1 RNA level. Twenty-four hr after A549 cells were transfected with siControl or sitRNase Z^L^, the cells were transfected without (mock) and with each of sgRNA, sgRM1, sgRM2, sgRM5 or sgRM6 (200 nM) and cultured further. After 96-hr culture, total cellular protein and RNA were prepared. (A) tRNase Z^L^ and β-actin protein levels were analyzed by Western blotting. (B−D) A MALAT1 RNA amount was measured by qRT-PCR, normalized against a β-actin mRNA amount, and expressed as a percentage relative to that of untreated cells. Data were from three biological replicates.

### Reduction in cell viability by suppression of the MALAT1 RNA level

Since it has been reported that suppression of the MALAT1 RNA level triggers apoptosis [17], we examined the sgRNAs sgRM1, sgRM2, sgRM5 (as a negative control) and sgRM6 for their ability to reduce cell viability. The effective sgRNAs sgRM1, sgRM2 and sgRM6 reduced A549 cell viability in 96 hr to 51, 58 and 67%, respectively (Fig 6).

**Fig 6.**
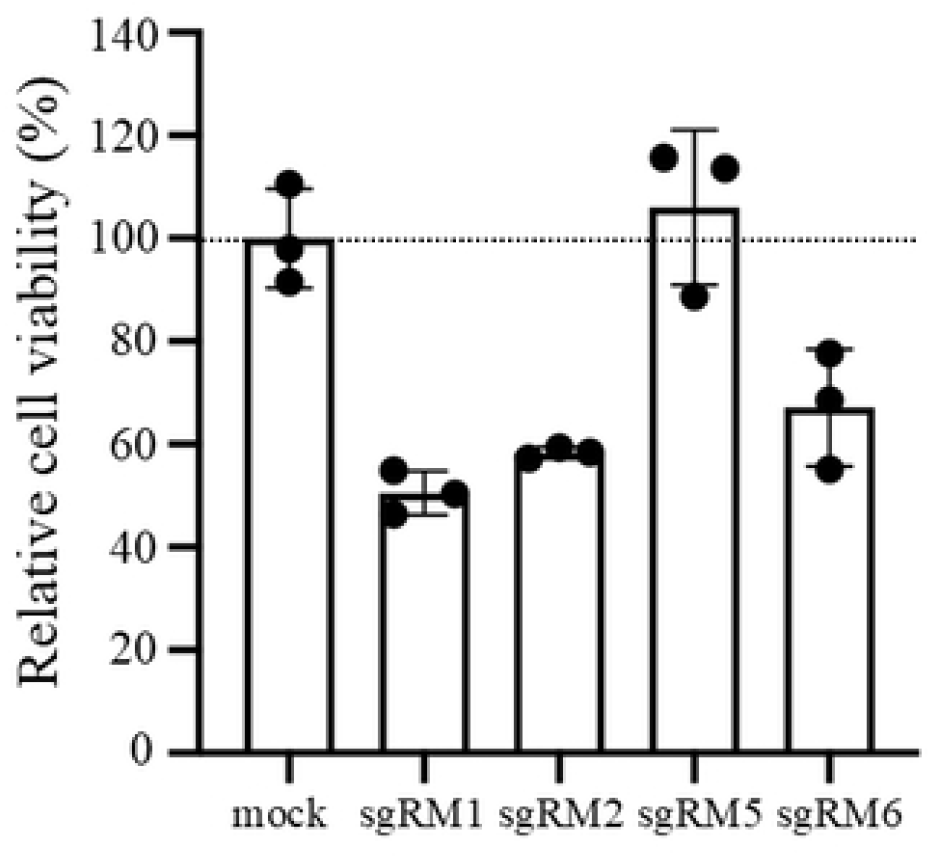
Cell viability assay. A549 cells were transfected without (mock) and with each sgRNA, sgRM1, sgRM2, sgRM5 or sgRM6 (200 nM), and after 96-hr culture, cell viability was measured. The cell viability is expressed as a percentage relative to that of untreated cells. Values are mean ± SD for three biological replicates.

## Discussion

We showed that the MALAT1 RNA level in A549 cells can be suppressed by transfecting them with 14-nt linear-type sgRNA, sgRM1, sgRM2 or sgRM6, and that the suppression is caused by TRUE gene silencing by presenting three pieces of evidence: the sgRNAs sgRM1, sgRM2 and sgRM6 can guide tRNase Z^L^ to cleave their own MALAT1 targets *in vitro*; a 14-nt linear-type sgRNA encapsulated into liposome can be delivered to the nucleus where both the target MALAT1 RNA and tRNase Z^L^ exist; downregulation of tRNase Z^L^ attenuates the suppression effect of sgRM1, sgRM2 and sgRM6 on the MALAT1 RNA level. Furthermore, we demonstrated that the effective sgRNAs sgRM1, sgRM2 and sgRM6 reduce cell viability.

Even the effective sgRNAs were hardly able to reduce the MALAT1 RNA level in 24 and 48 hr after transfection, and their suppression effect appeared only in 72 and 96 hr (Fig 2). The reason for this delay may be because tRNase Z^L^ needed to work primarily to produce mature tRNAs by removing 3′-trailers from pre-tRNAs in the relatively rapidly growing cells (corresponding to the cells 24 and 48 hr after transfection). In the slowly growing or quiescent cells (corresponding to the cells 72 and 94 hr after transfection), where tRNA 3′-processing would not be urgent, tRNase Z^L^ may become available to cleave target/sgRNA complexes even though the affinity of tRNase Z^L^ to the RNA complexes would be weaker than that to pre-tRNAs [22,23].

In contrast to sgRM1, the suppression effect of sgRM2 and sgRM6 did not show dose dependency over 200 nM (Fig 2B). Since it is well known that RNA, even relatively small one, can form various alternative conformations [24,25], the ∼10,500-nt long MALAT1 RNA will form many different conformations in the nucleus. The lack of dose dependency for sgRM2 and sgRM6 may be because a subset of MALAT1 RNA molecules form conformations in which binding sites of sgRM2 and/or sgRM6 are too tightly folded to access. The almost ineffective sgRNAs sgRM3, sgRM4 and sgRM5 (Fig 2A) would not be able to bind to most of the conformers of MALAT1 RNA, whereas sgRM1 may be able to bind to the MALAT1 RNA in its most of the alternative conformations.

In spite of the overwhelming reduction in the tRNase Z^L^ level, the sgRNAs sgRM1, sgRM2 and sgRM6 still showed their suppression effect on the MALAT1 RNA level, albeit different degree (Fig 5). The reduced amount of tRNase Z^L^ may be sufficient to cleave a certain amount of MALAT1 RNA molecules, or binding of the sgRNAs to MALAT1 may change its stable conformations to those susceptible to nuclease degradation regardless of the presence of tRNase Z^L^. Downregulation of tRNase Z^L^ slightly increased the MALAT1 level that was barely suppressed by sgRM5 (Fig 5B-D), suggesting that some conformers of MALAT1 RNA can be cleaved by sgRM5-guided tRNase Z^L^.

If we knew every conformation of a long RNA molecule in the cells, we could choose the best target site in the RNA. This might be realized in the future with the aid of artificial intelligence and/or quantum computers [26,27]. These technologies would also be used to predict off-target binding sites of sgRNAs in a transcriptome, and to avoid the off-target binding in designing sgRNA is very important especially in clinical application.

## Author Contributions

**Conceptualization:** Masayuki Takahashi, Masayuki Nashimoto.

**Data curation:** Masayuki Takahashi, Masayuki Nashimoto.

**Formal analysis:** Masayuki Takahashi, Masayuki Nashimoto.

**Funding acquisition:** Masayuki Nashimoto.

**Investigation:** Masayuki Takahashi.

**Project administration:** Masayuki Nashimoto.

**Resources:** Masayuki Takahashi, Masayuki Nashimoto.

**Supervision:** Masayuki Nashimoto.

**Validation:** Masayuki Takahashi, Masayuki Nashimoto.

**Visualization:** Masayuki Takahashi.

**Writing – original draft:** Masayuki Takahashi, Masayuki Nashimoto.

**Writing – review & editing:** Masayuki Nashimoto.

## Data Availability Statement

*All relevant data are within the manuscript*.

## Funding

*This work was supported by grant number 156 (to MN) from Takahashi Industrial and Economic Research Foundation. The funders had no role in study design, data collection and analysis, decision to publish, or preparation of the manuscript*.

## Competing interests

*The author MN is an advisor of Veritas In Silico Inc*., *and owns stock of the company*.

## Notes

### Competing Interest Statement

I have read the journal's policy and the authors of this manuscript have the following competing interests: The author MN is an advisor of Veritas In Silico Inc., and owns stock of the company.

## References

1. Nashimoto M, Kominami R, Nishi S, Mishima Y. A novel spermidine-dependent endoribonuclease activity caused by RNA-protein complex in mouse FM3A cell extracts. Biochem Biophys Res Commun. 1991 May 15;176(3):1163–9. doi: 10.1016/0006-291x(91)90407-x. PMID: 2039502.

2. Nashimoto M, Sakai M, Nishi S. Transfer RNA lacking its 3′ terminus is required for spermidine-dependent ribonuclease 65 activity in mouse FM3A cell extracts. Biochem Biophys Res Commun. 1991 Aug 15;178(3):1247–52. doi: 10.1016/0006-291x(91)91027-a. PMID: 1872844.

3. Nashimoto M. TRUE Gene Silencing. Int J Mol Sci. 2022 May 11;23(10):5387. doi: 10.3390/ijms23105387. PMID: 35628198; PMCID: PMC9141469.

4. Scherer L, Rossi JJ. Therapeutic potential of RNA-mediated control of gene expression: options and designs. In: Morris KV, editor. RNA and the regulation of gene expression: a hidden layer of complexity. Norfolk: Caister Academic Press Inc.; 2008. p. 201–226.

5. Thompson DM, Parker R. Stressing out over tRNA cleavage. Cell. 2009 Jul 23;138(2):215–9. doi: 10.1016/j.cell.2009.07.001. PMID: 19632169.

6. Shivarov V. TRUE gene silencing for hematologic malignancies. Leuk Res. 2014 Jul;38(7):729. doi: 10.1016/j.leukres.2014.04.014. Epub 2014 May 9. PMID: 24875835.

7. Takahashi M, Elbarbary RA, Nakashima A, Abe M, Watanabe N, Narita M, et al. A naked RNA heptamer targeting the human Bcl-2 mRNA induces apoptosis of HL60 leukemia cells. Cancer Lett. 2013 Jan 28;328(2):362–8. doi: 10.1016/j.canlet.2012.10.016. Epub 2012 Oct 22. PMID: 23092557.

8. Watanabe N, Narita M, Yamahira A, Taniguchi T, Furukawa T, Yoshida T, et al. Induction of apoptosis of leukemic cells by TRUE gene silencing using small guide RNAs targeting the WT1 mRNA. Leuk Res. 2013 May;37(5):580–5. doi: 10.1016/j.leukres.2013.01.015. Epub 2013 Feb 10. PMID: 23403166.

9. Takahashi M, Elbarbary RA, Watanabe N, Goto A, Kamiya D, Watabe Y, et al. Screening of a heptamer-type sgRNA library for potential therapeutic agents against hematological malignancies. Leuk Res. 2014 Jul;38(7):808–15. doi: 10.1016/j.leukres.2014.03.021. Epub 2014 Apr 5. PMID: 24768135.

10. Haino A, Ishikawa T, Seki M, Nashimoto M. TRUE Gene Silencing: Screening of a Heptamer-type Small Guide RNA Library for Potential Cancer Therapeutic Agents. J Vis Exp. 2016 Jun 2;(112):53879. doi: 10.3791/53879. PMID: 27285342; PMCID: PMC4927757.

11. Ishikawa T, Haino A, Ichiyanagi T, Seki M, Nashimoto M. Evaluation of double heptamer-type sgRNA as a potential therapeutic agent against multiple myeloma. Blood Cells Mol Dis. 2019 Nov;79:102341. doi: 10.1016/j.bcmd.2019.102341. Epub 2019 Jun 12. PMID: 31226499.

12. Ishikawa T, Haino A, Ichiyanagi T, Takahashi M, Seki M, Nashimoto M. Heptamer-type small guide RNA that can shift macrophages toward the M1 state. Blood Cells Mol Dis. 2021 Feb;86:102503. doi: 10.1016/j.bcmd.2020.102503. Epub 2020 Sep 7. PMID: 32920464.

13. Tamura M, Nashimoto C, Miyake N, Daikuhara Y, Ochi K, Nashimoto M. Intracellular mRNA cleavage by 3′ tRNase under the direction of 2′-O-methyl RNA heptamers. Nucleic Acids Res. 2003 Aug 1;31(15):4354–60. doi: 10.1093/nar/gkg641. PMID: 12888494; PMCID: PMC169917.

14. Takahashi M, Seki M, Nashimoto M. A naked antisense oligonucleotide with phosphorothioate linkages is taken up intracellularly more efficiently but functions less effectively. Biochem Biophys Res Commun. 2021 Oct 8;573:140–144. doi: 10.1016/j.bbrc.2021.08.035. Epub 2021 Aug 13. PMID: 34411896.

15. Arun G, Aggarwal D, Spector DL. MALAT1 Long Non-Coding RNA: Functional Implications. Noncoding RNA. 2020 Jun 3;6(2):22. doi: 10.3390/ncrna6020022. PMID: 32503170; PMCID: PMC7344863.

16. Gutschner T, Hämmerle M, Eissmann M, Hsu J, Kim Y, Hung G, et al. The noncoding RNA MALAT1 is a critical regulator of the metastasis phenotype of lung cancer cells. Cancer Res. 2013 Feb 1;73(3):1180–9. doi: 10.1158/0008-5472.CAN-12-2850. Epub 2012 Dec 14. PMID: 23243023; PMCID: PMC3589741.

17. Amodio N, Stamato MA, Juli G, Morelli E, Fulciniti M, Manzoni M, et al. Drugging the lncRNA MALAT1 via LNA gapmeR ASO inhibits gene expression of proteasome subunits and triggers anti-multiple myeloma activity. Leukemia. 2018 Sep;32(9):1948–1957. doi: 10.1038/s41375-018-0067-3. Epub 2018 Feb 22. PMID: 29487387; PMCID: PMC6127082.

18. Giard DJ, Aaronson SA, Todaro GJ, Arnstein P, Kersey JH, Dosik H, et al. In vitro cultivation of human tumors: establishment of cell lines derived from a series of solid tumors. J Natl Cancer Inst. 1973 Nov;51(5):1417–23. doi: 10.1093/jnci/51.5.1417. PMID: 4357758.

19. Elbarbary RA, Takaku H, Uchiumi N, Tamiya H, Abe M, Takahashi M, et al. Modulation of gene expression by human cytosolic tRNase Z(L) through 5′-half-tRNA. PLoS One. 2009 Jun 15;4(6):e5908. doi: 10.1371/journal.pone.0005908. PMID: 19526060; PMCID: PMC2691602.

20. Rossmanith W. Localization of human RNase Z isoforms: dual nuclear/mitochondrial targeting of the ELAC2 gene product by alternative translation initiation. PLoS One. 2011 Apr 29;6(4):e19152. doi: 10.1371/journal.pone.0019152. PMID: 21559454; PMCID: PMC3084753.

21. Brzezniak LK, Bijata M, Szczesny RJ, Stepien PP. Involvement of human ELAC2 gene product in 3′ end processing of mitochondrial tRNAs. RNA Biol. 2011 Jul-Aug;8(4):616–26. doi: 10.4161/rna.8.4.15393. Epub 2011 Jul 1. PMID: 21593607.

22. Nashimoto M, Wesemann DR, Geary S, Tamura M, Kaspar RL. Long 5′ leaders inhibit removal of a 3′ trailer from a precursor tRNA by mammalian tRNA 3′ processing endoribonuclease. Nucleic Acids Res. 1999 Jul 1;27(13):2770–6. doi: 10.1093/nar/27.13.2770. PMID: 10373595; PMCID: PMC148487.

23. Shibata HS, Takaku H, Takagi M, Nashimoto M. The T loop structure is dispensable for substrate recognition by tRNase ZL. J Biol Chem. 2005 Jun 10;280(23):22326–34. doi: 10.1074/jbc.M502048200. Epub 2005 Apr 11. PMID: 15824113.

24. Uhlenbeck OC. Keeping RNA happy. RNA. 1995 Mar;1(1):4-6. PMID: 7489487; PMCID: PMC1369058.

25. Nashimoto M. Correct folding of a ribozyme induced by nonspecific macromolecules. Eur J Biochem. 2000 May;267(9):2738–45. doi: 10.1046/j.1432-1327.2000.01294.x. PMID: 10785397.

26. Townshend RJL, Eismann S, Watkins AM, Rangan R, Karelina M, Das R, et al. Geometric deep learning of RNA structure. Science. 2021 Aug 27;373(6558):1047–1051. doi: 10.1126/science.abe5650. Erratum in: Science. 2023 Jan 27;379(6630):eadg6616. doi: 10.1126/science.adg6616. PMID: 34446608; PMCID: PMC9829186.

27. Fox DM, MacDermaid CM, Schreij AMA, Zwierzyna M, Walker RC. RNA folding using quantum computers. PLoS Comput Biol. 2022 Apr 11;18(4):e1010032. doi: 10.1371/journal.pcbi.1010032. PMID: 35404931; PMCID: PMC9022793.

